# Genomic dissection and mutation-specific target discovery for breast cancer *PIK3CA* hotspot mutations

**DOI:** 10.1101/2024.01.03.574067

**Authors:** Adam X. Miranda, Justin Kemp, Brad Davidson, Sara Erika Bellomo, Verda Agan, Alexandra Manoni, Caterina Marchiò, Sarah Croessmann, Ben H. Park, Emily Hodges

## Abstract

**Background:** Recent advancements in high-throughput genomics and targeted therapies have provided tremendous potential to identify and therapeutically target distinct mutations associated with cancers. However, to date the majority of targeted therapies are used to treat all functional mutations within the same gene, regardless of affected codon or phenotype.

**Results:** In this study, we developed a functional genomic analysis workflow with a unique isogenic cell line panel bearing two distinct hotspot *PIK3CA* mutations, E545K and H1047R, to accurately identify targetable differences between mutations within the same gene. We performed RNA-seq and ATAC-seq and identified distinct transcriptomic and epigenomic differences associated with each *PIK3CA* hotspot mutation. We used this data to curate a select CRISPR knock out screen to identify mutation-specific gene pathway vulnerabilities. These data revealed AREG as a E545K-preferential target that was further validated through *in vitro* analysis and publicly available patient databases.

**Conclusions:** Using our multi-modal genomics framework, we discover distinct differences in genomic regulation between *PIK3CA* hotspot mutations, suggesting the *PIK3CA* mutations have different regulatory effects on the function and downstream signaling of the PI3K complex. Our results demonstrate the potential to rapidly uncover mutation specific molecular targets, specifically AREG and a proximal gene regulatory region, that may provide clinically relevant therapeutic targets. The methods outlined provide investigators with an integrative strategy to identify mutation-specific targets for the treatment of other oncogenic mutations in an isogenic system.

## BACKGROUND

In the past few decades, significant strides in precision medicine and the advancement of targeted therapies have led to personalized treatment options and improved outcomes for patients with cancer, while limiting off-target toxicities. However, response to treatment still varies widely and the ability to better identify patients that would benefit from targeted therapies remains complex [1]. For the most part, current clinical practices regard mutations within the same gene as clinically equivalent despite distinct molecular differences, creating a significant obstacle in the implementation of targeted therapies [2, 3]. *PIK3CA*, which encodes the p110α subunit of phosphoinositide 3-kinase (PI3K), is the most commonly mutated gene in breast cancer and is responsible for regulating a diverse range of cellular functions including cell proliferation and survival [4-6]. *PIK3CA* has two distinct and highly prevalent hotspot mutations, E545K and H1047R, which occur in the helical and kinase domains, respectively, and have been shown to have distinct molecular changes and sensitivity to targeted therapeutics [5-9]. Despite these differences, current clinical application of PI3Kinase inhibitors in breast cancer do not distinguish between different mutations or between normal and mutated PI3K. This results in significant issues of toxicity, often times leading to dose reduction or discontinuation of the drug [2, 10-13]. To date, there is a distinct unmet need in the treatment of cancer, to accurately identify and understand molecular differences between mutations to effectively target cancer cells, improve selectivity, and decrease off-target effects.

Recent advancements in genomics technology and the affordability of generating high throughput genomics data have allowed researchers to begin to better understand the nuanced differences between mutations within the same gene. Furthermore, bioinformatic efforts have begun integrating transcriptomic and epigenomic data to better understand distinct molecular differences among unique mutational profiles from cancer patients. However, due to the significant mutational variability among individual cancers, as well as tumor heterogeneity and clonality, attributing observed differences to a single mutation has proven difficult. To better understand the molecular differences of *PIK3CA* hotspot mutations, our group has developed an integrative discovery platform to better identify key differences induced by different *PIK3CA* hotspot mutations in an isogenic human breast epithelial cell line panel [14]. The utilization of an isogenic mutation panel allows comparisons of the individual *PIK3CA* mutations under the expression of the endogenous promoter in near isolation, allowing for the identification of potential mutation-specific and mutation-preferential therapeutic targets.

The discovery platform presented here integrates RNA-seq, an assay for transposase-accessible chromatin with sequencing (ATAC-seq), and a select CRISPR knockout (KO) screen to uniquely identify distinct molecular targets attributed to either the *PIK3CA* E545K or H1047R mutations within a well-controlled model. RNA-seq allows for the identification and quantification of genes and pathways with altered expression due to the presence of either mutation [15]. ATAC-seq measures chromatin accessibility and can identify putative gene regulatory elements to provide additional insight into how regulation of genes and binding activities of transcription factors differ between two mutations within the same gene [16-18]. In our framework, data from these two assays are used to tailor a CRISPR screen that can accurately confirm genes with high essentiality in either mutant cell line; in doing so, we identify potential mutation-specific targets for treatment [19-21]. Combined application of these assays provides improved understanding of differences in cell function induced by distinct hotspot mutants as well as providing potential means of mutation-preferential inhibition.

Herein we describe a systematic approach (**Figure 1**) to identify potential mutation-preferential therapeutic targets. The utilization of an isogenic mammary epithelial cell model allows for the direct attribution of differences to specific mutations. This in return should improve selectivity of targeted therapies and decrease off-target effects. Our goal is to create a framework with “plug and play” accessibility for the evaluation of other hotspot mutants across cancer types using isogenic cell line models and to provide a foundation for future studies to identify a candidate list to maximize the potential for therapeutic benefit.

**Figure 1.**
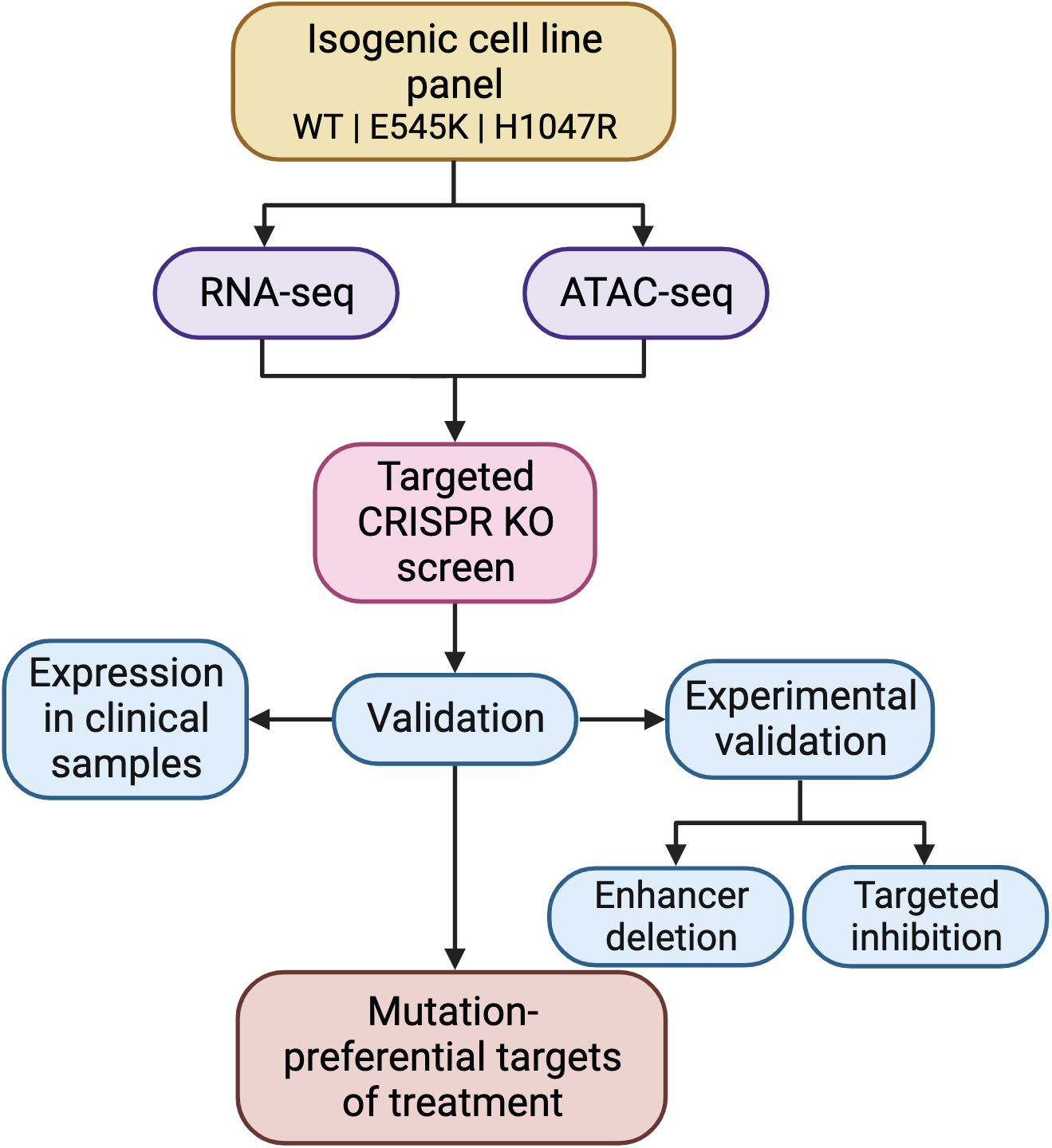
Discovery platform identifies mutation-preferential gene targets from isogenic cell line models. Flowchart breaking down the process of identifying selective gene targets from an isogenic cell line model.

## RESULTS

### RNA sequencing uncovers distinct transcriptional profiles and differential regulation of key cancer pathways in E545K and H1047R *PIK3CA* mutant cells

To evaluate differences in the transcriptomes of cells harboring the *PIK3CA* hotspot mutations E545K and H1047R, we performed RNA-seq on a panel of isogenically modified nontumorigenic breast epithelial MCF-10A cell lines harboring the respective mutations. RNA-seq identified 1271 genes with differential expression between the two mutant cell lines (**Figure 2A and 2B**) [22]. A complete summary of the differentially expressed genes (DEGs) can be found in **Table S1** and are displayed in **Figure 2B**. Interestingly, hierarchical clustering revealed the gene expression patterns of the E545K cell line shared greater similarity with the WT parental cell line than the H1047R mutant cell line (**Figure 2A**).

**Figure 2.**
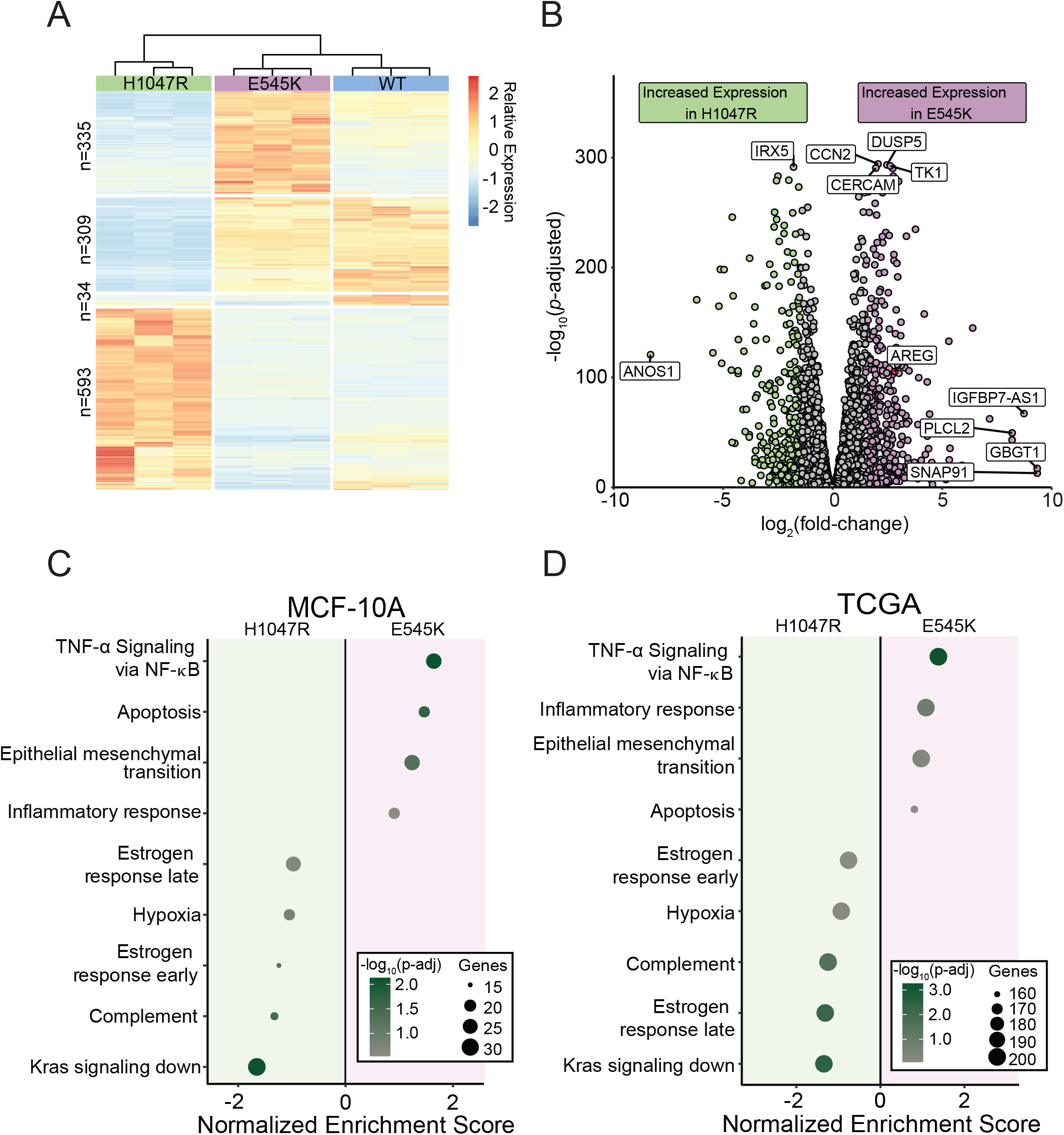
RNA-seq captures distinct gene expression differences induced by *PIK3CA* hotspot mutations in isogenic cell line models which are reflected in TCGA patient samples. (A) Heatmap of normalized counts for 1271 differentially expressed genes. Hierarchical clustering of these genes reveals that E545K cells bear more similarity to WT than H1047R. (B) Volcano plot of differential expression between mutant isogenic cell lines. Differentially expressed genes (DEGs) were defined by the criteria: fold change > |1.5|, *P*_adj_ < 0.05. (C, D) Dot plot showing results from GSEA pathway enrichment analyses using the hallmark gene sets for (C) MCF-10A DEGs and (D) expression data from TCGA-BRCA samples. Pathways shown in panel D are those that are significantly enriched, shared and concordant with those identified as significant in the MCF-10A cell line.

Gene set enrichment analysis using the MSigDB Hallmark pathway collection was performed to identify patterns of shared function across DEGs [23]. Multiple uniquely enriched pathways were identified for each mutant (**Figure 2C**). Pathways related to cell cycle and proliferation, as well as epithelial-mesenchymal transition genes, were enriched in the E545K cells, while estrogen response pathways and K-ras associated genes were enriched in the H1047R cells. These differences in gene expression patterns suggest distinct modes of tumorigenic activity between the different mutants, despite being treated as clinically equivalent. Using patient data from The Cancer Genome Atlas (TCGA) Breast Cancer (BRCA) data set, RNA-seq samples from tumors bearing each *PIK3CA* mutation confirmed observations made within our isogenic MCF-10A panel (**Figure 2D**). Of the 14 pathways found to be significantly enriched in our panel, all were confirmed to be significantly enriched in differentially expressed gene sets from corresponding TCGA mutant samples. These findings demonstrate single amino acid substitutions in the same gene can have wide-ranging and distinct disruption of gene expression, which translate directly to expression differences observed in clinical samples.

### PIK3CA mutants demonstrate unique differences in chromatin accessibility and gene regulation

Considering the gene expression changes observed with RNA-seq, we performed ATAC-seq to identify genomic regions with altered regulatory landscapes, which may contribute to changes in gene expression. In addition to the identification of dynamic regions of chromatin accessibility (ChrAcc), differences in transcription factor (TF) binding activities were also estimated from Tn5 cut-site profiles. Comparing accessibility profiles between E545K and H1047R mutants identified 8672 differentially accessible regions. We performed unsupervised clustering to define 4 distinct groups of accessibility patterns (**Figure 3A**). Two distinct groups, designated as E545K-preferred and H1047R-preferred, represented putative regulatory loci with increased accessibility in either E545K or H1047R mutant cells, respectively. In addition to providing insight into gene regulatory mechanisms, ChrAcc provides insight into *cis*-regulatory elements including enhancers. Within both the E545K-preferred and H1047R-preferred region clusters, over 50% of the regions fall within intronic and distal intergenic sequences (**Figure 3B**). These results suggest that a significant amount of chromatin remodeling that is driven by the different *PIK3CA* mutations occurs at noncoding enhancer elements that can bind TFs and influence gene expression [24].

**Figure 3.**
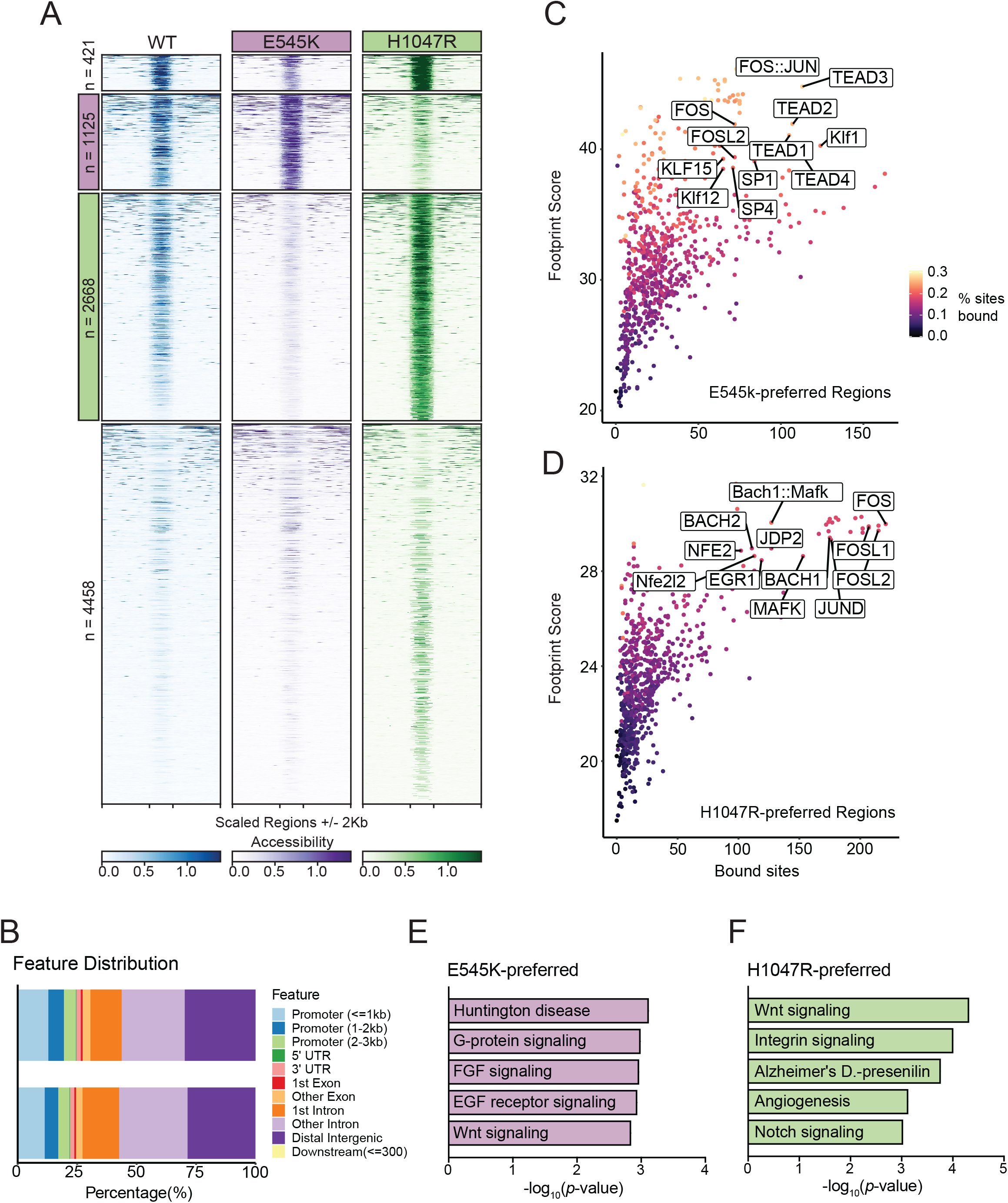
ATAC-Seq identifies mutation-specific gene regulatory mechanisms near genes of key pathways. (A) Heatmap of accessibility at 8672 peaks exhibiting differential accessibility between the mutant isogenic cell lines. Peaks are divided based on k-means clustering. The second cluster highlighted in pink has been designated as E545K-preferred. The third cluster highlighted in green has been designated as H1047R-preferred (B) Distribution of genomic feature annotations of regions within the E545K-preferred and H1047R-preferred clusters. (C) Scatter plots of TOBIAS transcription factor footprinting of accessibility in the E545K-preferred regions. (D) Scatter plots of TOBIAS transcription factor footprinting of accessibility in the H1047R-preferred regions. (E) Bar plot of pathway enrichment from the PANTHER pathway database analysis performed on genes uniquely annotated to the E545K-preferred cluster. (F) Bar plot of pathway enrichment from the PANTHER pathway database analysis performed on genes uniquely annotated to the H1047R-preferred cluster regions.

Indeed, TF motif analysis revealed that each accessibility cluster is enriched for distinct families of TF binding sites identified from the JASPAR database [25, 26] (**Figure S1**). In the E545K-preferred cluster, strong enrichment for the hormone receptor transcription factors ARE and PGR were observed as well as the TEAD transcription factor family. The TEAD TF family has been shown to have a strong association with canonical PI3K/AKT signaling and can promote epithelial to mesenchymal transition [27, 28]. In the H1047R-preferred cluster, there was increased enrichment of AP-1 family TFs. AP-1 family TFs have been shown to interact with chromatin remodelers and promote a proliferative gene expression program [29, 30], and have also been associated with signaling through the MAPK cascade [31].

While TF motif analysis informs which sequences are enriched within ChrAcc regions, it does not predict TF occupancy. To better understand differential TF binding activities, we performed TF footprinting using TOBIAS, which uses Tn5 cut-site profiles to identify differences in proteins bound at TF binding motifs [17]. Our results show high levels of TEAD TF binding in E545K-preferred regions (**Figure 3C**) and high levels of AP-1 binding (FOS, FOSL1, FOSL2, JUND) in H1047R-preferred regions (**Figure 3D**). These results are consistent with the motif enrichment results and point to activity of the TEAD and AP-1 TF families as key regulators of differential gene expression between the *PIK3CA* hotspot mutants.

Nearest neighbor gene annotation using GREAT was used for gene ontology analysis to identify genes uniquely associated with either mutation-preferred accessibility cluster and analyzed for pathway enrichment using Enrichr [32-35]. Using the PANTHER database, we identified enrichment of distinct pathways promoted by either *PIK3CA* mutants [36, 37]. Within the E545K-preferred cluster regions, unique enrichment of multiple growth factor receptor signaling pathways was observed and is likely due to changes in PI3Kα signaling induced by the E545K mutant cells (**Figure 3E**) [4]. Interestingly, enrichment from the H1047R-preferred cluster regions showed enrichment for both the Notch and Wnt signaling pathways. Both of these pathways are associated with the promotion of tumor growth in breast cancers, but neither are canonically associated with increased activity of PI3Kα **(Figure 3F**) [38, 39]. This suggests H1047R mutant cells may drive alternative proliferative cell signaling outside of canonical PI3K signaling. These gene ontology results are consistent with the observed TF enrichment and footprinting between clusters, and provide additional context to the differential gene expression observed from RNA-seq. The differences in ChrAcc demonstrate distinct differences in genomic regulation between *PIK3CA* mutations and suggest the *PIK3CA* mutations have different effects on the function and downstream signaling of the PI3K complex.

### Select CRISPR-Cas9 knockout screen identifies genes with mutation-specific essentiality

A key advantage of our isogenic cell line model is the ability to compare both mutants to the unmodified parental cell line. Therefore, a CRISPR KO screen could accurately identify essential genes specific to *PIK3CA* mutations, but not *PIK3CA* WT cells, and may provide a list of promising therapeutic targets with limited off-target effects in normal cells. Performing a whole genome CRISPR screen can be both time and resource intensive. To circumvent these limitations, we used the data generated from both the previously performed RNA-seq and ATAC-seq assays to curate a select list of genes to investigate within our CRISPR KO screen. Analysis of RNA-seq data identified 616 DEGs, 160 for E545K mutants and 456 for H1047R mutants, with significantly upregulated expression in a mutant cell line relative to the parental (**Figure 4A and Figure 4B**). Among the 9672 combined mutant-specific genes that annotate to regions of differential chromatin accessibility, 312 were identified as DEGs (**Figure 4C**).

**Figure 4.**
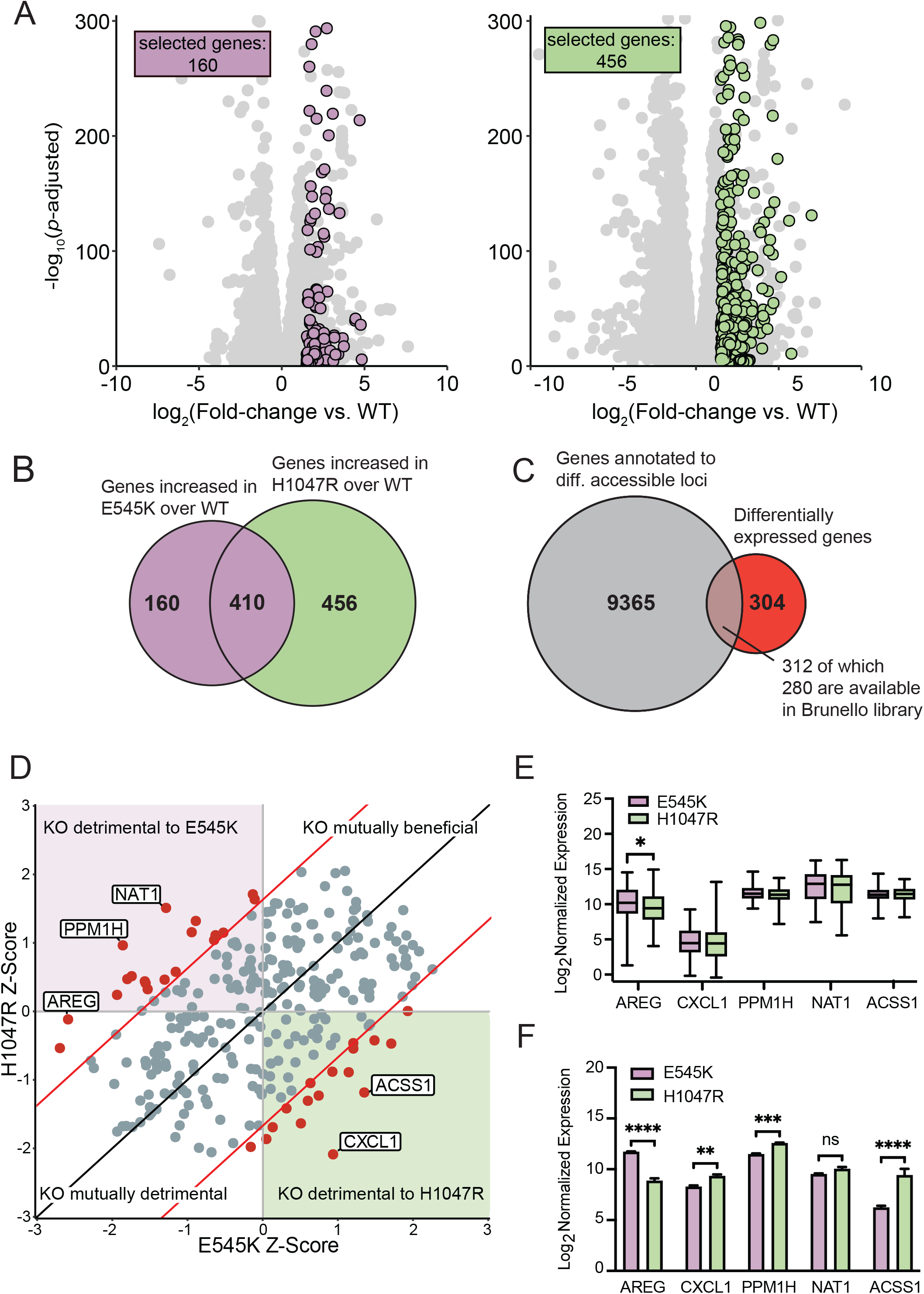
Genes with altered expression and nearby chromatin accessibility were selected for a CRISPR KO screen to identify gene targets with mutation-specific essentiality. (A) Scatter plots of gene expression in mutant cells relative to parental cells. Unique genes meeting the fold change and significance threshold (fold change > |1.5|, *P*_adj_ < 0.05) shown in color. (B) Euler plot showing the overlap of genes with increased expression in cells with either mutation compared to the parental cell line. (C) Euler plot showing the overlap between genes annotated to regions of differential accessibility with those exhibiting increased expression in a mutant cell line compared to the parental cell line. (E) Box plots showing expression of the top 5 hits from the CRISPR screen in TCGA-BRCA samples. AREG shows significant difference in expression between samples bearing either of the hotspot mutations. (F) Bar plots showing the expression of the top 5 hits from the CRISPR screen in the MCF-10A RNA-seq samples.

Among the 312 selected genes, 280 genes with targeting single guide RNAs (sgRNAs) were available from the Brunello full genome library for a select CRISPR KO screen [40] (**Table S2**). Using the MAGeCK software package, Z-score differences between each of the mutant cell lines were compared to the parental cell line for each gene [41] (**Figure 4D**). From this analysis, we identified 36 genes with a Z-score difference greater than a significance threshold of 1.65, which corresponds to a confidence interval of 95% (**Table S3**). When knocked out, these genes specifically disrupt the survival of either mutant cell line with minimal disruption to the parental line (**Table S3 and Figure 4D**). The top five genes (NAT1, PPM1H, AREG, ACSS1, CXCL1) with the greatest differences in Z-scores were evaluated in the *PIK3CA* mutant breast cancer samples from TCGA (**Figure 4E**). Of the top 5 genes, AREG was the only gene with significant differential expression between the *PIK3CA* mutations. Samples with E545K mutations demonstrated a significant increase in expression when compared to the H1047R, recapitulating the differential expression of AREG observed in our isogenic panel (**Figure 4F**). This association was independently confirmed utilizing tumor samples from the Molecular Taxonomy of Breast Cancer International Consortium (METABRIC) database, with an increased difference within the luminal B subtype (**Figure S2**) [42]. Clinical confirmation of a unique molecular target identified from the select CRISPR screen emphasizes the translational potential of hits identified from isogenic mutant cell lines analyzed within our strategy.

### Disruption of differentially accessible locus identified by ATAC-seq exhibits regulatory function over AREG expression

Selection criteria for inclusion of the ATAC-seq data in the CRISPR screen required identification of differentially accessible peaks between the two mutants. The accessible peak annotated to AREG (chr4:74435384-74435596 locus) was identified as significantly more accessible in the E545K mutant cells and exhibits many qualities of a gene regulatory region **(Figure 5A**). A previous study using CTCF ChIA-PET in MCF-10A cells published by ENCODE (ENCSR403ZYJ) showed that this locus interacts with the promoter of the AREG gene and could influence expression [43, 44]. Furthermore, the Genotype-Tissue Expression project (GTEx) identifies 5 different AREG expression quantitative trait loci (eQTLs) single nucleotide polymorphisms (SNPs) within 1kb of this region (**Table S4**) and these SNPs have been shown to influence the expression of AREG in multiple tissue types (**Figure 5A**). Specifically, the rs28570600 SNP (gold square, **Figure 5A**) has previously been shown to be significantly associated with breast cancer susceptibility [45].

**Figure 5.**
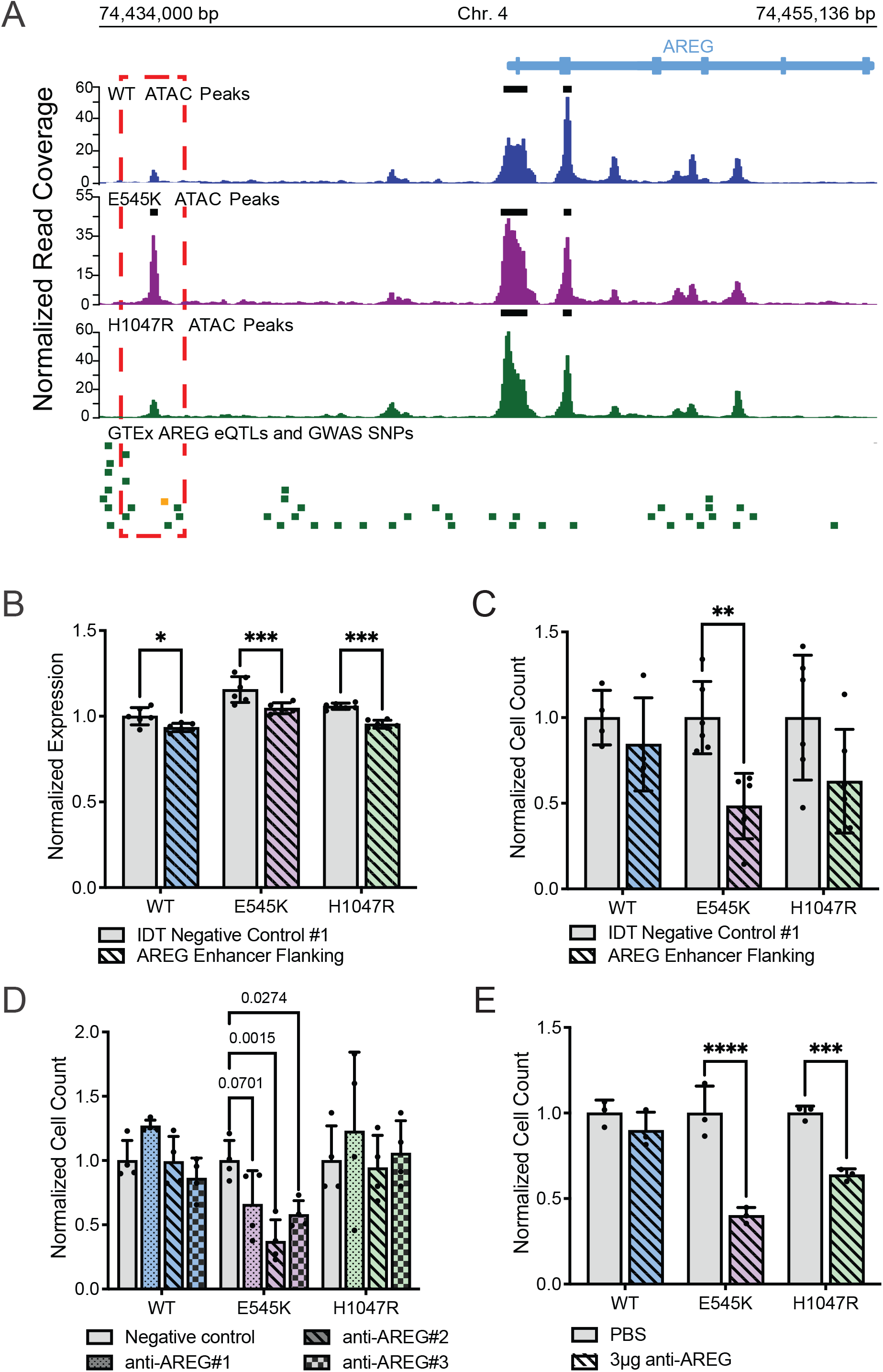
E545K mutant cells exhibit specific dependence on *AREG*, which is regulated by nearby accessibility peak/putative enhancer. (A) Genomic tracks showing the ATAC-seq data across the isogenic cell lines alongside key SNPs at the *AREG* gene locus. The red box highlights the differentially accessible region annotated to the *AREG* gene. The green bars designate GTEx *AREG* eQTLs. The gold bar designates a GWAS SNP associated with breast cancer. (B) Bar plot of *AREG* expression following CRISPR-mediated deletion of the putative *AREG* enhancer. (C) Bar plot of cell counts following CRISPR-mediated deletion of the putative *AREG* enhancer. (D) Bar plot showing differences in survival/proliferation following inhibition of *AREG* expression using siRNA (Oligos on **Table S5**, expression of *AREG* shown in **Figure S3**). (E) Bar plots showing differences in survival/proliferation following inhibition of AREG using a neutralizing antibody (R&D Systems, MAB262-SP).

To investigate the function of this peak, the region was deleted using CRISPR-Cas9 and a pair of sgRNAs targeting the discussed locus upstream of the AREG TSS (oligo sequences in **Table S5**). Loss of this enhancer region significantly reduced AREG expression in all cell lines in the isogenic model (**Figure 5B**). There was also an observed decline in the proliferation/survivability of the mutant cell lines, however only significant within the E545K cells. (**Figure 5C**). These assays demonstrate the capability of our pipeline to identify regulatory regions that may themselves provide targets for mutation-specific treatment of *PIK3CA* mutant disease.

### *PIK3CA* mutant cells exhibit specific dependency on AREG

To confirm the role of AREG expression on cell survival, short interfering RNA (siRNAs) were used to inhibit AREG in the isogenic panel (**Figure S3**). With an siRNA knockdown, our goal was to assess the effect of a reduction of AREG expression without complete loss of expression in the system. Consistent with observations from the CRISPR KO screen, reduction of AREG expression significantly disrupted the survival and proliferation of E545K mutant cells, while exhibiting no significant changes in WT or H1047R mutant cells (**Figure 5D**). To validate this observation, cells were treated with a neutralizing antibody that disrupts the extracellular signaling of AREG. Both mutant cell lines exhibited sensitivity to AREG perturbation, while WT cells showed no significant change (**Figure 5E**). The extracellular nature of AREG makes it a particularly attractive target as inhibitors of AREG would not necessarily need to penetrate the cell membrane to be effective. This could simplify drug design and reduce potential off target toxicity [46, 47]. Taken together, these results demonstrate the utility of the pipeline to identify potential molecular targets as an option for mutation-preferential therapeutic strategy. The *in vitro* and translational confirmation of our findings demonstrate the power of our model to accurately identify actionable gene targets and gene regulatory regions with high selectivity for mutant cells and minimal impact to WT cells.

## DISCUSSION

Increased availability and advancements in multi-omics technology have begun to revolutionize translational research to better understand the interplay of molecular changes and provide new opportunities for targeted therapies. However, integration and implementation of multi-omics data for identifying new molecular targets for therapeutic development remains underutilized in the cancer setting. This study presents an analytical pipeline employing an isogenic mutant panel to better understand and uniquely identify the molecular differences between mutations within the same gene. Traditionally, most cancer-associated mutations have been clinically evaluated and treated as a monolithic group with variable success. More contemporary targeted therapies such as inhibitors specific to mutant KRAS G12C, for example, highlight the success and feasibility of developing mutation specific inhibitors [48]. To improve upon this current paradigm, our workflow takes advantage of a model of isogenically incorporated mutations in a genetically stable background, integrating both RNA-seq and ATAC-seq data to design a uniquely tailored CRISPR KO screen enabling the detection of mutation-selective targets. The utilization of a mutant model that incorporates an isogenic background provides a system to identify a candidate list that demonstrates mutation-preferential gene regulatory dependencies. Furthermore, the accessibility of CRISPR-Cas9 gene editing systems makes the development of isogenic models for cancer-associated mutations a relatively fast and straightforward process and can be scaled for a variety of mutations across tumor types. In addition to identifying mutation-preferential molecular targets, our comprehensive process paired with the isogenic panel can identify and characterize potential enhancer regions with mutation-specific activity that may offer alternative targets for treatment. These putative enhancers have affinity for distinct TF families that result in unique expression profiles and may be exploited as therapeutic vulnerabilities. The true utility of our process is in the identification of potential targets. Hits from our analyses still require additional validation to determine their effects beyond the CRISPR KO screen and ultimately their translation for potential clinical impact.

Evidence for the applicability of our workflow in breast cancer was used to analyze the two most common *PIK3CA* mutations in breast cancer to identify distinct molecular differences that impact downstream signaling, chromatin accessibility, and gene expression. RNA-seq and ATAC-seq analysis identified the disruption of epithelial-mesenchymal transition associated genes in E545K mutant cells and the MAPK cascade in H1047R mutant cell. Integration of this information to create a uniquely tailored, focused CRISPR screen allowed us to identify AREG as an E545K-specific exploitable molecular difference in a highly efficient manner. AREG has been previously and independently established as a signaling molecule required for the growth of PIK3CA-mutant breast cancer cells [49]. Independent identification of this molecular target utilizing our pipeline demonstrates its immediate biological application. Furthermore, we were able to confirm translational applicability through retrospective analyses of publicly available patient data. While our experiments were performed in a single isogenic cell line model, these clinical findings suggest the applicability of our results to actual breast cancer patients. Taken together, this study provides a framework for the independent evaluation of oncogenic hotspot mutations from a functional genomics perspective. This implies that in the era of patient-specific treatment and pharmacogenomics, our process may allow for the discovery of new targets and improved personalized medicine with the potential for increased specificity and decreased toxicity.

## CONCLUSIONS

This work highlights the utility of integrating multiomics data collected from an isogenic mutant model to better identify molecular targets for therapy. With the increased accessibility to genome editing technology and services, our pipeline can provide investigators with a clear method for studying specific mutants in other cancer cell line models. Our workflow was able to identify AREG as an E545K-preferential molecular target, which was confirmed through *in vitro* assays and retrospective analyses of patient data, highlighting the potential clinical utility of our work.

## MATERIALS AND METHODS

### Cell Culture

MCF-10A parental cell lines were purchased from American Type Culture Collection (ATCC). MCF-10A cell line knock-ins were generated as previously described [14]. All cell lines were grown in 5% CO_2_ at 37°C with 1% Penicillin/Streptomycin in respective media conditions. Parental MCF-10A cell lines were cultured in DMEM/F12 (1:1) supplemented with 5% horse serum, 20 ng/ml epidermal growth factor (EGF), 10 µg/ml insulin (Roche), 0.5 µg/mL hydrocortisone (Sigma), and 100 ng/ml cholera toxin (Sigma). Knock-in cell lines were maintained in MCF-10A media in the absence of EGF. For all sequencing assays, cells were transferred to assay media 24 hours prior to sample collection. Assay media contains phenol red-free DMEM/F12 (1:1) supplemented with 1% charcoal-dextran stripped FBS (Fisher), 0.2 ng/ml EGF, 10 µg/ml insulin, 0.5 µg/mL hydrocortisone, and 100 ng/ml cholera toxin.

### RNA-Seq

RNA was isolated and prepared using the Qiagen RNeasy kit. Libraries were prepared by the Vanderbilt Technologies for Advanced Genomics (VANTAGE) Core using the Illumina Ribo-Zero Plus rRNA Depletion Kit. Each library was sequenced on an Illumina NovaSeq, PE150, at a requested depth of 50 million reads. All code and the specific parameters used in all data analyses can be found at: (https://github.com/adamxmiranda/PIK3CA). All sequencing library reads were trimmed of adapters and assessed for quality using the Trim Galore! (version 0.4.0) Wrapper of Cutadapt and FastQC[50, 51]. Trimmed reads were mapped to the human genome assembly hg38 using the Spliced Transcripts Alignment to a Reference (STAR) aligner (version 2.5.4b) [52]. Mapped reads were sorted and filtered for a mapping quality score over 30 using the SAMtools package (version 1.5) [53]. Reads were counted to gene transcripts using featureCounts (version 2.0.0) to version 32 of the GENCODE transcripts [54, 55]. A summary of the sequencing preprocessing can be found in **Table S6**. Differential gene expression was identified between conditions using the DESeq2 package [22]. Pathway analysis was performed using the fgsea package on the Hallmark gene set from MsigDB [23, 56].

### ATAC-seq

Nuclei were isolated and ATAC-seq libraries were prepared using previously published methods [16, 57]. Libraries were sequenced by the VANTAGE Core on an Illumina NovaSeq PE150, at a requested depth of 50 million reads. Reads from the ATAC-seq libraries were trimmed using the same process described in the RNA-seq section. All code and specific parameters used in all data analyses can be found at: (https://github.com/adamxmiranda/PIK3CA). Trimmed reads were mapped to the human genome assembly hg38 using the BBTools (version 38.69) package and Burrows-Wheeler Aligner (version 0.7.17) [19, 58]. Quality filtering was performed on the mapped reads using SAMtools [53]. A summary of the sequencing preprocessing can be found in **Table S6**. Peaks of accessibility were called using Genrich (version 0.6.1) and differential accessibility was determined using DESeq2 (version 1.34.0) [22]. Accessible regions were clustered using k-means clustering. Gene annotation and pathway enrichment was performed using GREAT (version 4.0.4) [32, 33]. The gene annotation parameters used for GREAT were the default parameters of 5kb upstream of the transcriptional start site (TSS), 1kb downstream of the TSS, or up to 1000kb in either direction for distal regions. Pathway enrichment was performed on uniquely annotated genes using the Enrichr web browser tool and the PANTHER database [34-37]. Motif enrichment and transcription factor (TF) footprinting were performed using HOMER (version 4.10) and TOBIAS (version 0.13.3), respectively, to identify TF potentially binding to identified accessible peak clusters [17, 26].

### CRISPR KO Screen

A modified CRISPR screen was performed with a select cohort of gene targets selected based on two criteria: 1. genes exhibited significantly increased RNA expression (log2 fold change greater than 1.5 and p-adjusted value less than 0.05) specifically in one of the *PIK3CA*-mutant cell lines compared to wild type, and 2. genes were annotated to regions that demonstrated significantly increased accessibility in either mutant cell line. Differential accessibility was assessed using DESeq2 and nearest neighbor annotation for these regions was performed using ChIPseeker (version 1.30.3) with the default annotation conditions of +/- 3kb from the TSS [22, 59]. Based on this selection criteria, 312 genes were selected, for which 280 had guides available in the Brunello whole genome single guide RNA (sgRNA) library. The Brunello whole genome sgRNA library was modified for these 280 genes and prepared by the Vanderbilt Functional Genomics core in the lentiCRISPRv2 plasmid background (Full list of guides **Table S2**) [19, 60].

MCF-10A cells were cultured in the maintenance media conditions at a density of 500,000 cells/well in a 6-well plate. 24 hours after seeding, cells were infected with viral supernatant in maintenance media containing 5 μg/mL polybrene. 24 hours post-infection cells were placed in selection media containing 1 μg/mL of puromycin and maintained for two weeks. Following two weeks of selection, libraries were prepared and sequenced following the protocol described in Sanjana *et al* [60]. Libraries were sequenced by the VANTAGE core and analysis was performed using the maximum likelihood estimation (MLE) algorithm within the MAGeCK software package(version 0.5.9.5) [41, 61].

### Deletion of putative AREG Enhancer

Two sgRNAs were designed targeting the locus, chr4:74435384-74435596, which is annotated to the AREG gene as well as nearby AREG eQTLs identified from the GTEx Portal [62-64] (**Table S4**). The guide RNAs were purchased as sgRNAs from IDT with custom targeting sequences (full guide sequences in **Table S5**). Cells were plated in respective maintenance media conditions at a density of 50,000 cells/well in a 12-well plate. Transfection mixtures were prepared and added to each of the cell lines according to the guidelines described in the Lipofectamine CRISPRMAX Transfection Reagent kit (Invitrogen, CMAX00001) using the provided reagents and the Alt-R S.p. HiFi Cas9 Nuclease V3 (IDT, 1081060) in Opti-MEM Reduced Serum Media (ThermoFisher, 31985062). Separate mixtures were prepared containing either a 1:1 mixture of the designed AREG enhancer flanking guides or a control mixture of Alt-R® CRISPR-Cas9 Negative Control crRNA #1 (IDT, 1072544) duplexed to Alt-R® CRISPR-Cas9 tracrRNA (IDT, 1072532). DNA and RNA were collected from separate experimental replicates 24 hours after transfection (at least 4 technical replicates were prepared for each cell line in each treatment condition). Cell counts and viability were measured 72 hours following transfection using a Vi-CELL BLU cell viability analyzer (Beckman Coulter).

DNA was collected using the Wizard SV 96 Genomic DNA Purification System (Promega, A2370). PCR was performed using Phusion High-Fidelity PCR Master Mix (NEB, M0531) and the enhancer deletion validation primer set (**Table S5**). PCR mix and thermocycler conditions were set according to the Phusion Master Mix protocol provided by NEB. PCR products were measured and visualized using a D5000 ScreenTape System (Agilent, 5067) (**Figure S4**).

RNA was isolated from cells using the RNeasy Plus Mini Kit (Qiagen, 74134). RNA was converted to cDNA using the iScript cDNA Synthesis Kit (Bio-rad, 1708890). qPCR was performed using the AREG and ACTB qPCR primer sets (**Table S5**) for each sample with the SYBR Green PCR Master Mix (Applied Biosystems, 4309155). Expression of AREG was calculated relative to the expression of housekeeping gene ACTB.

### Anti-AREG Antibody Assay

Cells were plated in their respective maintenance media conditions at a density of 50,000 cells/well in 12-well plates. 24 hours following seeding, cells were treated with 1, 3, or 5 µg of AREG neutralizing antibody (R&D Systems, MAB262-SP) or an equivalent volume of PBS. This experiment was performed with three replicates for each cell line and each treatment condition. Cells were counted and viability was measured 72 hours following treatment using a Vi-CELL BLU cell viability analyzer (Beckman Coulter).

### siRNA Assay

Cells were plated in their respective maintenance media at a density of 50,000 cells/well in a 12-well plate. 24 hours following seeding, cells were treated with three different commercially validated AREG targeting siRNA (Ambion, see oligo sequences on **Table S5**), a negative control siRNA (Invitrogen, 4390843), or a null transfection condition using the Lipofectamine RNAiMAX Transfection Reagent (Invitrogen, 13778100) at a concentration of 10 pmol siRNA per well. 24 hours post-transfection, RNA was prepared from cells using the Qiagen RNeasy kit. Four replicates were prepared from each cell line and each treatment condition. RNA was converted to cDNA using the iScript cDNA Synthesis Kit (Bio-rad, 1708890). qPCR was performed using the AREG and ACTB qPCR primer sets (**Table S5**) for each sample with the SYBR Green PCR Master Mix (Applied Biosystems, 4309155). Expression of AREG was calculated relative to the expression of housekeeping gene ACTB.

To assess the impact on survival and proliferation, cells were plated in their respective maintenance media at a density of 30,000 cells/well in a 24-well plate. Cells were treated with one of three different commercially validated AREG targeting siRNA (Ambion, see oligo sequences on **Table S5**) or a negative control siRNA (Invitrogen, 4390843) using the Lipofectamine RNAiMAX Transfection Reagent (Invitrogen, 13778100) at a concentration of 5 pmol siRNA per well. Cell counts and viability were measured 24 hours following treatment using a Vi-CELL BLU cell viability analyzer (Beckman Coulter). This experiment was performed with four replicates in each cell line and each treatment condition.

### Visualization and Figure Creation

Images and figures were generated using ggplot2 (version 3.4.1), plotgardener (1.4.1), deeptools (3.5.1), pheatmap (1.0.12), and graphpad Prism (Version 10) [65-68]. Schematic images and flow chart were created using Biorender.com.

## ABBREVIATIONS

PIK3CA: Phosphatidylinositol-4,5-Bisphosphate 3-Kinase Catalytic Subunit Alpha
PI3K: Phosphatidylinositol-4,5-Bisphosphate 3-Kinase
ATAC-seq: Assay for transposase-accessible chromatin with sequencing
CRISPR: Clustered Regularly Interspaced Palindromic Repeats
KO: Knock out
AREG: Amphiregulin
DEGs: Differentially expressed genes
ChrAcc: Chromatin accessibility
TF: Transcription factor
eQTL: Expression quantitative trait locus

## DECLARATIONS

### Ethics approval and consent to participate

Not applicable.

### Consent for publication

Not applicable.

### Availability of Data

Raw sequencing datasets are available through the Gene Expression Omnibus (GEO) with the accession number: GSE247822. Detailed workflows and all code used in data analysis can be found at: (https://github.com/adamxmiranda/PIK3CA).

### Competing Interests

CM reports personal consultancy fees from Bayer, Roche, AstraZeneca, Novartis, outside the scope of the present work.

### Funding

We are grateful for support of the project and the time invested in producing this manuscript by NIH awards [1R01GM147078-01 to E.H], Department of Defense Idea Award [W81XWH-20-1-0522 to E.H], American Cancer Society (ACS) Institutional Research Grant (#IRG-15-169-56), the Vanderbilt Stanley Cohen Innovation Fund and funds from the Vanderbilt Ingram Cancer Center. We would also like to acknowledge support from the Breast Cancer Research Foundation, the Susan G. Komen Foundation, and the NIH CA214494, CA194024 (B.H.P.). We would also like to thank and acknowledge the support of The Canney Foundation, the Sage Patient Advocates, the Marcie and Ellen Foundation, The Eddie and Sandy Garcia Foundation, the support of Amy and Barry Baker, the support of John and Donna Hall, and the Vanderbilt-Ingram Cancer Center support grant (NIH CA068485) and Breast Cancer SPORE (NIH CA098131).

### Authors’ Contributions

AXM, BHP and EH conceived of the project. AXM cultured the cells and performed the data analysis for the RNA-seq, ATAC-seq and CRISPR KO screen experiments. AXM performed the enhancer deletion and siRNA experiments. JK performed the anti-AREG antibody experiments. BD and AM assisted with cell culture. BD and VA contributed to experimental design. SEB and CM provided data and analysis of METABRIC RNA-seq samples. AXM, SC, BHP and EH wrote and edited the manuscript and figures. All authors reviewed and approved the final manuscript.

## Acknowledgements

The results shown here are in part based upon data generated by the TCGA Research Network: https://www.cancer.gov/tcga. Schematic images created with biorender.com.

The Genotype-Tissue Expression (GTEx) Project was supported by the Common Fund of the Office of the Director of the National Institutes of Health, and by NCI, NHGRI, NHLBI, NIDA, NIMH, and NINDS. The data used for the analyses described in this manuscript were obtained from: the GTEx Portal on 11/02/2022.

