## Supplementary material for "Genomic dissection and mutation-specific target discovery for breast cancer *PIK3CA* hotspot mutations": FigS1

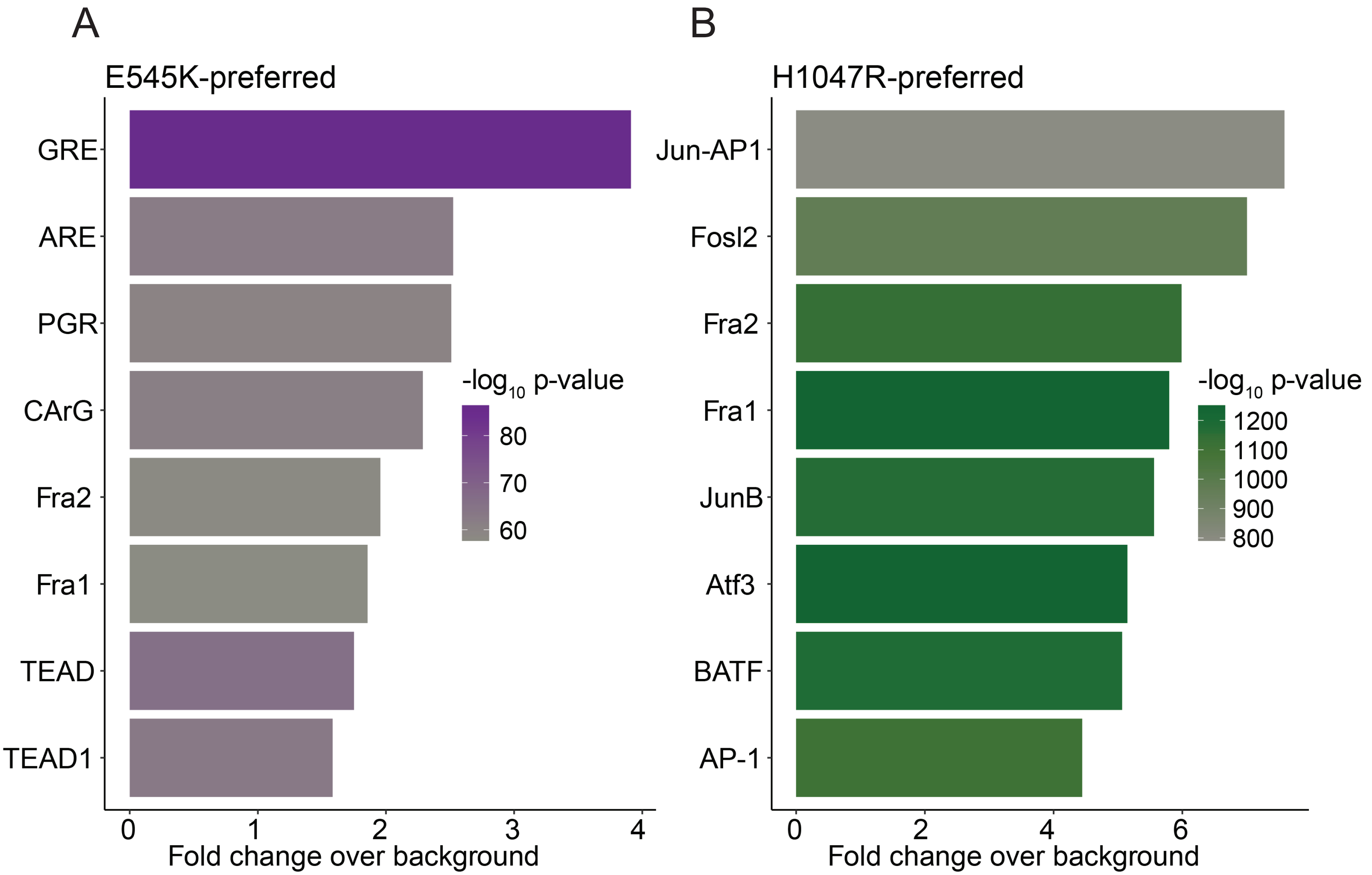


Figure S1.
Bar plots showing the results of HOMER TF motif enrichment analysis within the (A) E545K-preferred and (B) H1047R-preferred cluster regions.
