## Supplementary material for "Genomic dissection and mutation-specific target discovery for breast cancer *PIK3CA* hotspot mutations": FigS2

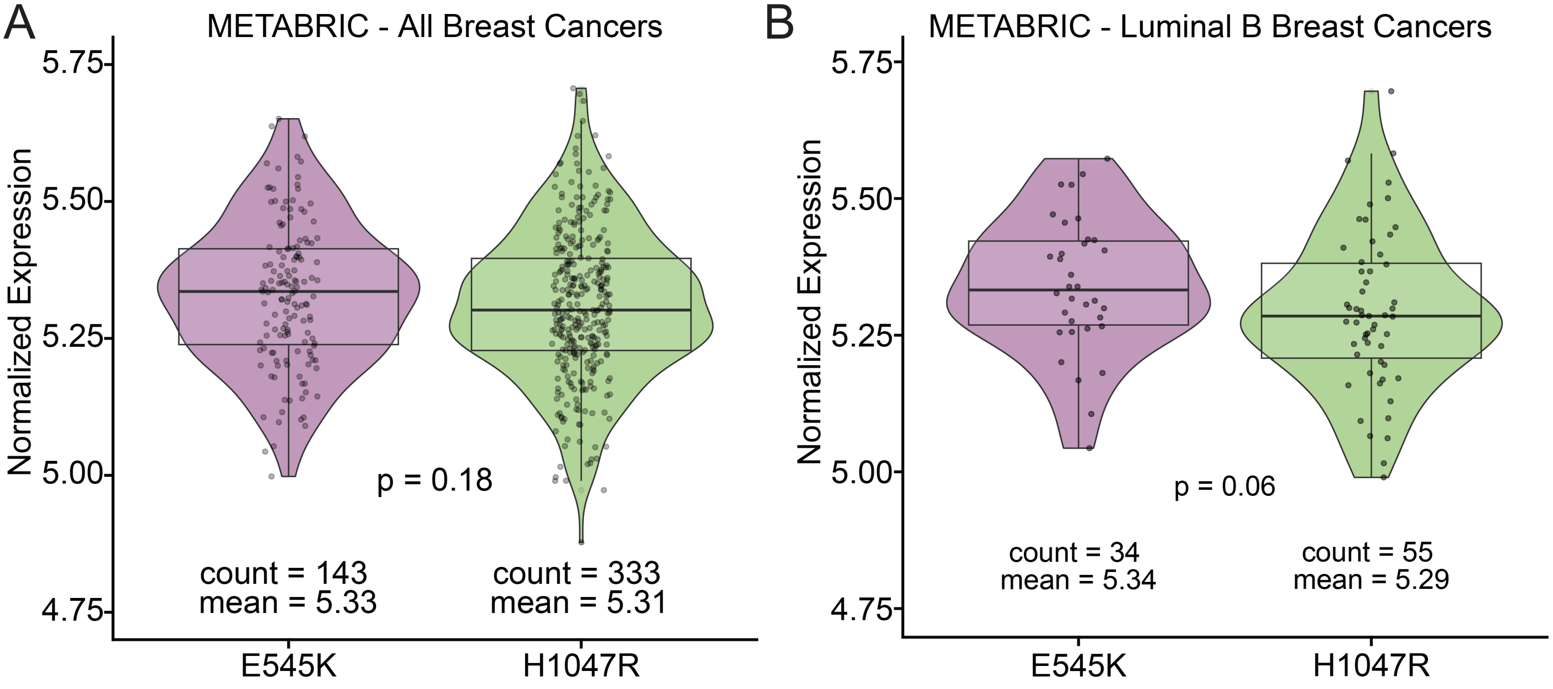


**Figure S2.**
Violin plots showing the expression of AREG from METABRIC samples with either PIK3CA hotspot mutation. (A) AREG expression in all samples. (B) AREG expression from samples specifically of the luminal B subtype.
