## Supplementary material for "Genomic dissection and mutation-specific target discovery for breast cancer *PIK3CA* hotspot mutations": FigS3

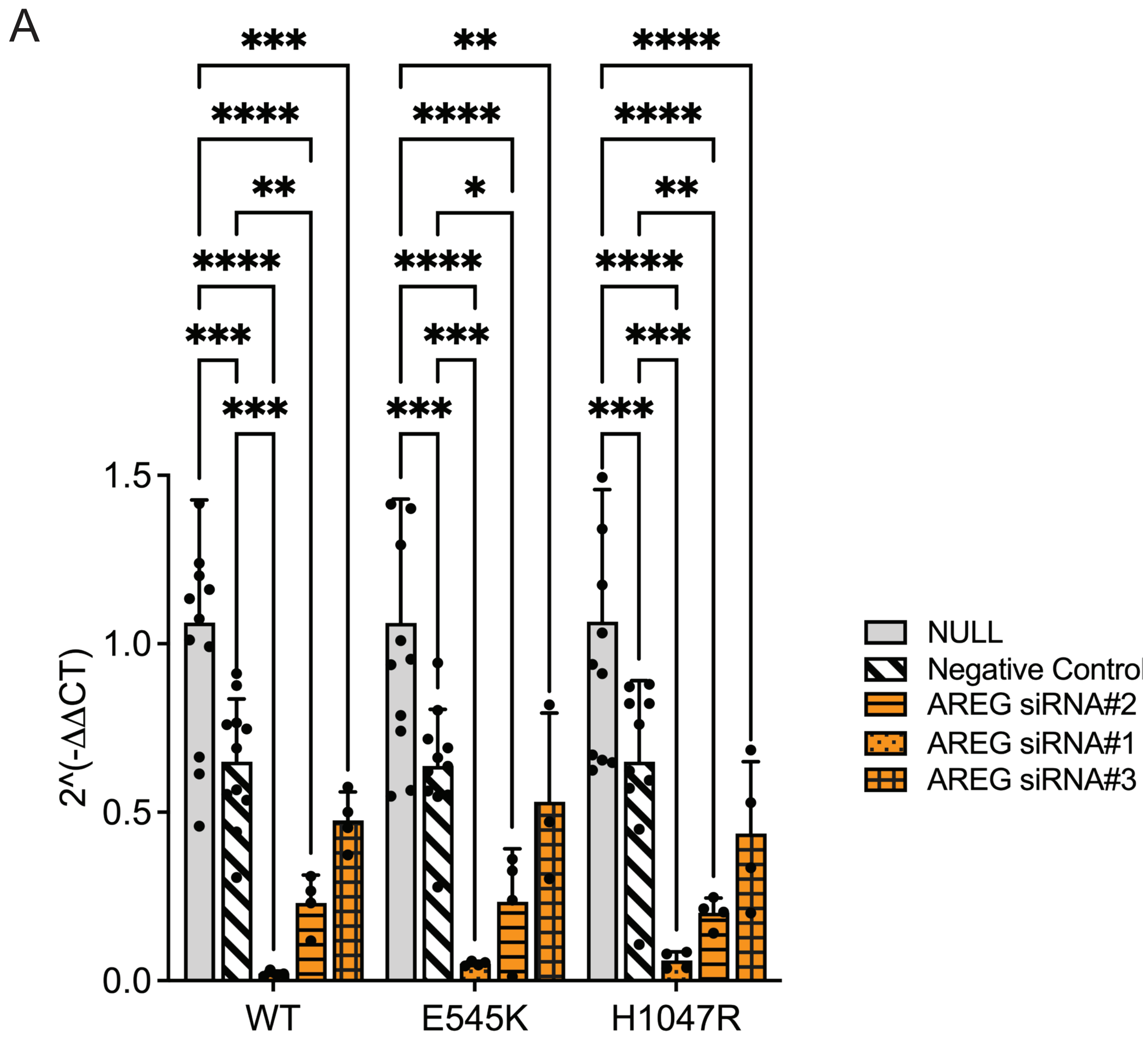


**Figure S3.**
(A) Bar plots showing the differences in the expression of AREG relative to ACTB following siRNA-mediated inhibition of AREG with each of the three AREG-targeting siRNA molecules listed in Table S5.
Outliers were removed using the ROUT method with a Q threshold of 1%.
