## Supplementary material for "Genomic dissection and mutation-specific target discovery for breast cancer *PIK3CA* hotspot mutations": FigS4

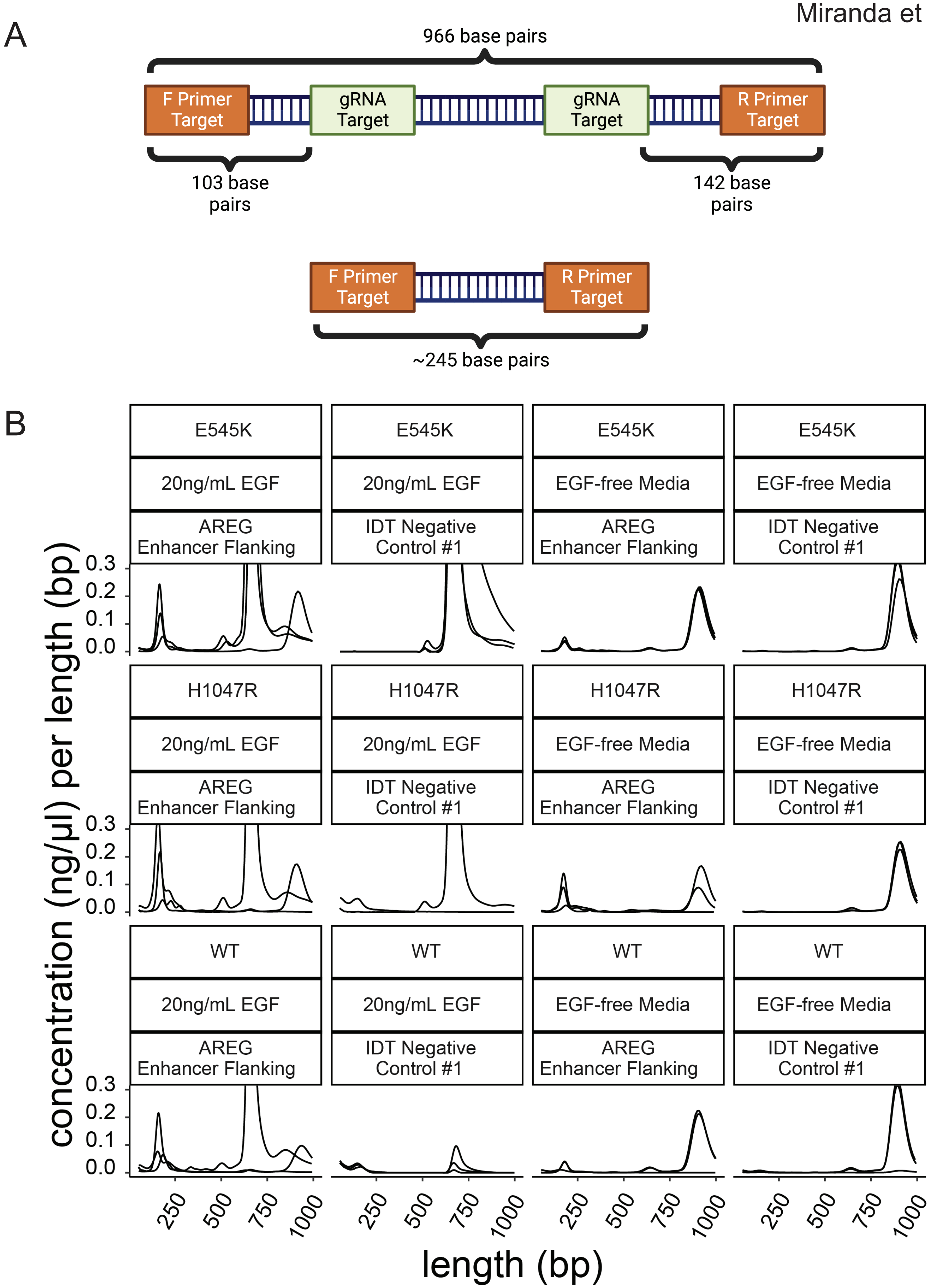


**Figure S4.**
Validation of enhancer region deletion. (A) Schematic showing the predicted PCR product sizes based on the deletion of the targeted region. A PCR product of 966 bp corresponds to the unmodified enhancer region. The PCR product of ~245bp signifies a deleted enhancer. (B) Reduced electropherograms from D5000 tapestation from 3 replicates of all cell lines and all media conditions. The presence of the ~250bp band is more prevalent in cells that received the AREG enhancer flanking guides compared to the negative control.
